# Enzymatic Degrading Chlorophenol Wastewater by Mixed Strains of Immobilized White Rot Fungi†

**DOI:** 10.1101/2024.09.09.611973

**Authors:** Hongyuan Liu, Xueyi Lou, Yeyao Shao, Zhichao Wang, Jiamin Xiao, Kai Cen, Dingyin Chen, Qiman Xia, Wenlong Xu, Fang Fang, Yasin Orooji, Peng Liu

## Abstract

To address chlorophenol wastewater pollution, immobilized mixed white rot fungi (WRF) strain microsphere was designed as a solid degradation agent, using *lignin peroxidase* (*LiP*), *manganese peroxidase* (*MnP*), and *laccase* (*Lac*) to degradating the wastewater. Considering the diverse physical and chemical properties of the fungal sphere, the immobilization agent formula is optimized and comprehensive environmental factor design response surface analysis are implemented to determine the delivery conditions. Consequently, the 2,4-DCP treatment rate and extracellular enzyme activity for a 1:1 encapsulation of *T. versicolor* and *P. sajor-caju* significantly outperform those of individual strains. Using polyvinyl alcohol (PVA), sodium alginate (SA), and biochar as carriers, with sodium dihydrogen phosphate solution as crosslinker and SiO2/zeolite as additives, immobilizing mixed bacteria yielded a high-quality solid agent. This achieved a 99.33% 2,4-DCP degradation rate over 96 hours, with optimal dosage, pH, and initial 2,4-DCP concentration at 11.5 g/L, 5.5, and 40 mg/L. The degradation of 2,4-DCP by WRF selectively removes adjacent chlorine atoms to produce 4-CP, enhancing the dechlorination efficiency.

**Graphical Abstract:** 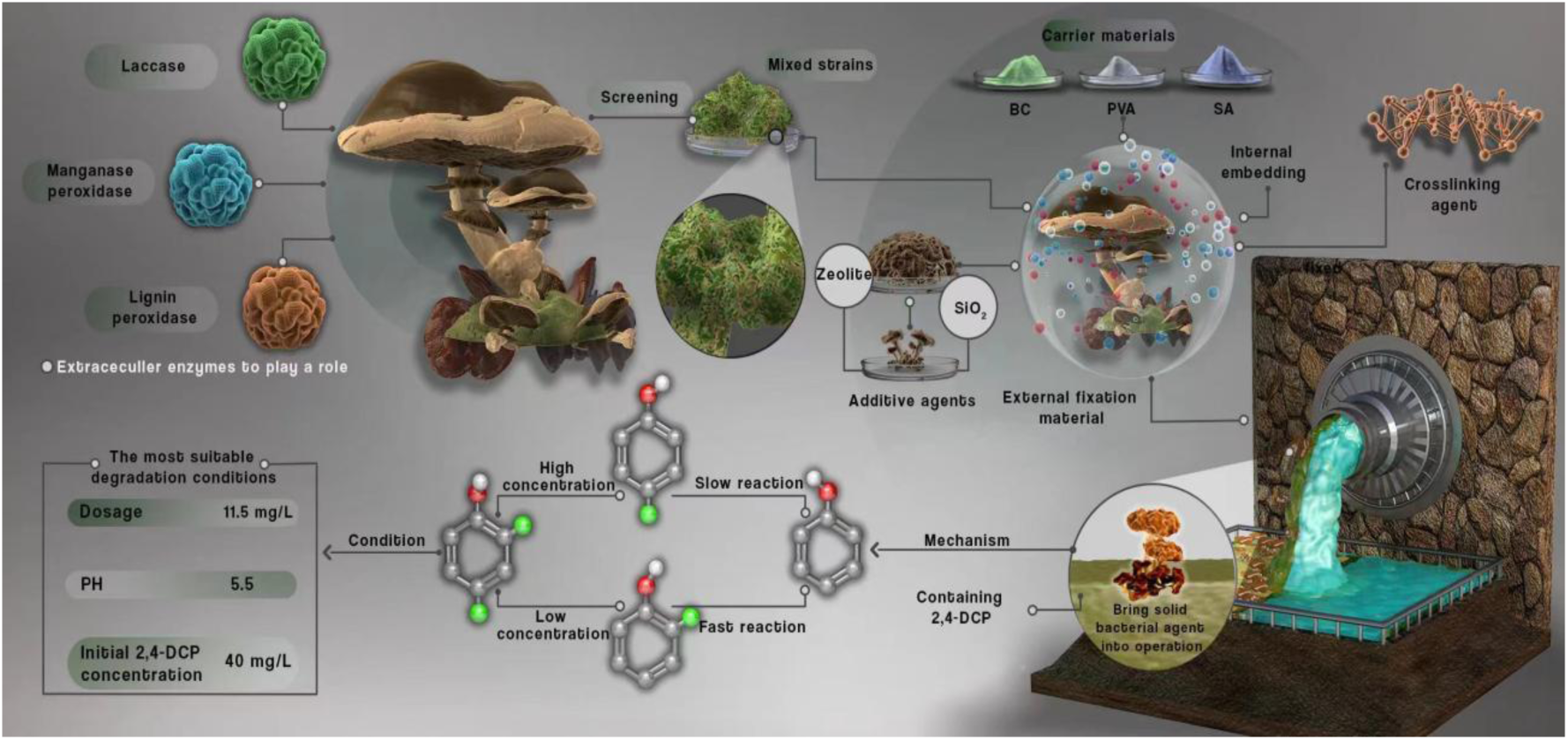

## 1 Introduction

2,4-dichlorophenol (2,4-DCP) stands out as the primary intermediate in the degradation pathway of the herbicide 2,4-dichlorophenoxyacetic acid (Liu et al., 2020). It induces cells to generate reactive oxygen species (ROS), disrupting the bilayer structure of cell membrane phospholipids, and exhibits potent carcinogenic and mutagenic effects on organisms (Hu et al., 2024). Classified as a priority pollutant by the US Environmental Protection Agency, addressing 2,4-DCP pollution is imperative to human society (He et al., 2021). While physicochemical methods for removing 2,4-DCP from sewage entail high energy consumption and generate additional pollutants, biological methods offer the advantages of being cost-effective, efficient, and environmentally sustainable. Therefore, to investigate efficient approaches to mitigate 2,4-DCP pollution is important, with biological methods being particularly promising in the pursuit of environmental sustainability (Zhang et al., 2021).

White Rot Fungi (WRF), a group of filamentous fungi renowned for their ability to utilize and decompose plant lignin (Ma et al., 2021), leading to white wood decay, are recognized for their extensive degradation of organic compounds, including 2,4-dichlorophenol (2,4-DCP), thus providing its potential candidate in addressing environmental pollutions. Moreover, WRF species are extensively employed in the degradation of synthetic pesticides, polycyclic aromatic hydrocarbons (PAHs), and polychlorinated biphenyls (PCBs), owing to their unique capability to decompose pollutants and fix them in mineral form (Harguindeguy et al., 2024). Generally, species such as *Phanerochaete chrysosporium* (Kato et al., 2024), *Pleurotus ostreatus* (Zhuo & Fan, 2021), *Trametes versicolor* (Tan et al., 2023), and *Pleurotus sajor-caju* (del Cerro et al., 2021) are notable examples of WRF known for their production of key degrading enzymes, including *lignin peroxidase* (*LiP*) (Wang et al., 2023a), *manganese peroxidase* (*MnP*) (Wu et al., 2023), and *laccase* (*Lac*). Through a sequence of free radical chain reactions, these enzymes facilitate the conversion of chlorophenol into environmentally benign inorganic chloride, CO_2_, and H_2_O.

However, it worth noting that each enzyme activity is different in the separated strain species, this reveled that the single reaction is not capable to support a sequence reaction under the single enzyme activity (Tortella et al., 2008). The excessive intermediate products remain after single treatment. To remove the organic contaminants thoroughly, mixed cultivation effectively mitigates the adverse impacts of excessive intermediate products on fermented production and finds extensive applications in both production processes and pollution management. For example, Zhang et al. (2024) utilize a mixed fungi remediation process involving *Pleurotus dryinus* and *Trametes hirsuta*, they achieved high rates of phenol removal while also contributing to the elimination of other organic and heavy metal compounds. Similarly, Kaewlaoyoong et al. (2020) employed mixed white rot fungi (specifically *Pleurotus pulmonarius*) to degrade polychlorinated dibenzo-p-dioxins (PCDDs), dioxins, and furans in contaminated soil via solid-state fermentation, achieving an impressive removal rate of 96%.

Second, considering for the harsh environment of waste water, immobilized microbial technology offers several advantages. This technology involves fixing dominant microorganisms capable of degrading specific substances onto a carrier material, thereby effectively enhancing microbial density. This enhancement assists in improving the removal rate of pollutants and isolates direct contact between microorganisms and the polluted environment, alleviating the impact of environmental changes on water treatment efficacy and helps prevent the loss of dominant bacteria. Moreover, PVA-immobilized cells have proven effective in biodegrading various harmful contaminants, including aromatic compounds (Messinis et al., 2024), phenolic compounds (Palumbo et al., 2024), and crystal violet (Rahmatpour et al., 2024). Following the degradation process, microorganisms can absorb toxic ions or convert pollutants into harmless byproducts. The wrapping of immobilized materials prevents intermediate substances from being easily released into the environment until complete degradation occurs, ensuring that only small molecule substances are released.

Upon these concerns, herein, the 2,4-DCP degradation capabilities of four WRF combining immobilization with mixed bacteria fermentation technology are investigated. Through the intriguing selection of mixed strains, the reasonable collocation formula of the different enzymic catalytic properties of *LiP*, *MnP*, and *Lac* can be acquired. To strengthen the strains’ stability under ambient conditions, the immobilization technique was implemented to encapsulate WRF with PVA, SA, and biochar as embedding material, and the solid immobilized microspheres could be finally formed. To elucidate the advantage of the current degradation agent, response surface analysis is discussed to illustrate the optimal experimental conditions, and the corresponding dichlorination degradation mechanism of 2,4-DCP by mixed WRF strains is given. We believe this report could shed light on the further investigation of biological treatment of organic pollutants under a green and sustainable methodology without energy consumption.

## 2 Materials and methods

### 2.1 Materials and reagents

The *P. chrysosporium* (BKMF-1767), *T. versicolor* (ATCC 42530), *P. ostreatus* (HK 35), and *P. sajor-caju* (ATCC 32078) used in this study were obtained from the China Center for Type Culture Collection. WRF was maintained by subculturing on potato dextrose broth (PDB) slants at 4°C, and then introduced into potato dextrose agar (PDA) plates for 10 days at 37°C. As WRF mature, remove the spores, and dissolve them into sterile distilled water, and then adjusted to a concentration of 2.0×10^6^ CFU/mL by using spectrometer (X-8S, Shanghai, China).

2,4-DCP was purchased from Chinese Chemical Research Institute, 500 mg/L of 2,4-DCP stock solution was prepared and stored at 4°C. By diluting the stock solutions, different concentrations of 2,4-DCP were prepared. All other reagents used were of analytical grade and were purchased from **Shanghai Aladdin Biochemical Technology Co., Ltd.**

### 2.2 Preparation of PVA-immobilized WRF beads (PWBs)

The method of gel-making has been improved on Chen’s method (Chen et al., 2022). It was prepared by adding PVA (10%, w/v), SA (1%, w/v), silica (2%, w/v) and zeolite powder (1%, w/v) to a 0.9% sodium chloride solution and stirring slowly in an 80°C water bath. Dissolve NaH2PO4 in saturated boric acid solution to obtain 1.0 M sodium dihydrogen phosphate solution, which could be used as cross-linking agent autoclaved with gel solution at 111℃ for 30 min, then could be used after cooling to about 45°C.

The WRF spore suspension (20%, w/v) was added to the rice straw biochar (1.5%, w/v) and Tween-80 (0.05%, v/v), then add it to the gel solution and mixed well, placing in a shaker at 28°C for a period of 2 h. The biochar and colloid adsorbed by WRF were mixed and stirred. After mixing and stirring, the mixture was added dropwise to CaCl_2_ (2%, w/v) solution using 5 mL syringe and cured for 2 h and placed on a magnetic stirrer (MS3, Zhejiang, China). After curing, they were washed with sterile 0.9% NaCl and refrigerated at 4°C.

### 2.3 Determination of Chlorophenols

Add 50 mL liquid to the colorimetric tube, add 0.5 mL of ammonia water-ammonium chloride buffer solution with a pH of 10.0±0.2, use 1.0 mL of 4-aminoantipyrine solution as the chromogenic agent. Then add 1.0 mL of potassium ferricyanide solution as the oxidizing agent, mix thoroughly and seal for 10 minutes. Measure the absorbance value of the solution, using a colorimetric dish with an optical path of 20 mm at a wavelength of 510 nm, using phenol free water as a reference to obtain Chlorophenols in concentration.

### 2.4 Dechlorination of 2,4-DCP

Prior to measurement, each liquid sample was filtered through a 0.22 μm membrane filter. 2,4-DCP and its dechlorination intermediates 2-CP, 4-CP and phenol were measured using a high-Performance Liquid Chromatograph (HPLC, model: 1100, Agilent, USA) equipped with a 250 nm diode array detector (DAD) and the column (ZORBAX SB-C18, California, America) (Akerman-Sanchez & Rojas-Jimenez, 2021). The column temperature was 25°C, the flow rate was 1mL/min, the detector wavelength was 530 nm and the sample size was 20 μL (Banerjee et al., 2024). The mobile phase was methanol : acetate : water solution (70:1:29, v/v/v).

### 2.5 Microorganism and growth conditions

Stock cultures were maintained at 4℃ on malt extract agar slant. Suspensions of the mycelia was prepared in sterile distilled water. A turbidimeter (WGZ-200, Shanghai, China) was used to measure the fungal concentration. The concentration was adjusted to 2.0×106 CFU/mL. For inoculation of the liquid culture medium, aqueous suspensions of fungal spores were prepared. Kirk’s liquid culture medium (200 mL) was stored in a 500-mL Erlenmeyer flask. The flasks were incubated for 3 days at 37°C in an incubator at 150 rpm with 6×10^6^ spores (v/v=1.5%) (Dao et al., 2021).

### 2.6 Effect of pH, initial 2,4-DCP concentration

Different concentrations of 2,4-DCP were prepared at 20 mg/L, adjusting at the beginning of the experiment with 0.1 mol/L HNO_3_ or NaOH, and the effect of pH on the degradation rates was investigated over a pH range of 3.5 to 8.5. Initial 2,4-DCP concentrations were adjusted to 0, 5, 10, 20, 80, 100, 120, and 160 mg/L. The optimum initial concentration of 2,4-DCP and the effect of the initial concentration of 2,4-DCP on the removal of 2,4-DCP were investigated (Kots et al., 2023).

### 2.7 Enzymes activities assay

The oxidation of veratryl alcohol to veratraldehyde at 25°C by hydrogen peroxide was monitored at A310 nm to determine lignin peroxidase (*LiP*) activity in the extracellular medium (Kommedal et al., 2023). One unit of *LiP* activity was defined as the amount of enzyme required to produce 1 μmol of veratryl aldehyde per minute under the assay conditions (Zhang et al., 2020). The activity of manganese peroxidase (*MnP*) has been measured by monitoring the change in oxidation state from Mn^2+^ to Mn^3+^ at A240 nm (Yadav et al., 2023). The reaction started at a temperature of 25°C. The amount of enzyme required to produce 1 μmol Mn^3+^ from the oxidation of Mn^2+^ per minute was used as the unit of *MnP* activity. The spectrophotometric measurements were carried out using a UV-visible light spectrophotometer (Su et al., 2024).

### 2.8 Statistical analysis

The experimental test data were calculated by Excel 2016, one-way analysis of variance (ANOVA) by SPSS 26.0 software, followed by Ducan method for multiple comparisons (α=0.05), JMP Pro 16 software for mixed design, and selected the related experiments of response surface optimization in Design-Expert 8.0 software. Origin Pro 2022 software mapping and all data are presented as mean ± standard deviation. Each experiment and the index measurement were set three times in parallel.

## 3. Results

### 3.1 Enzyme activity of WRF and degradation rate of 2,4-DCP by single WRF bacteria

The degradation of organic pollutants by WRF is mainly dependent on the secretion of lignin modifying enzymes and their ability to degrade pollutants, and this enzyme complex is known for its high oxidation capacity and lack of specificity. It can be seen from the analysis of **Fig. 1** that there are significant differences in the enzyme activities of *MnP*, *Lac* and *LiP* enzymes of the four strains. The enzyme activity of *MnP* enzyme in *P. ostreatus* is significantly higher than that of *Lac* enzyme and *LiP* enzyme, while *MnP* enzyme in *P. ostreatus* attacks phenolic hydroxyl group through peroxidation and produces reactive intermediate (peroxide free radical).

**Fig. 1.**
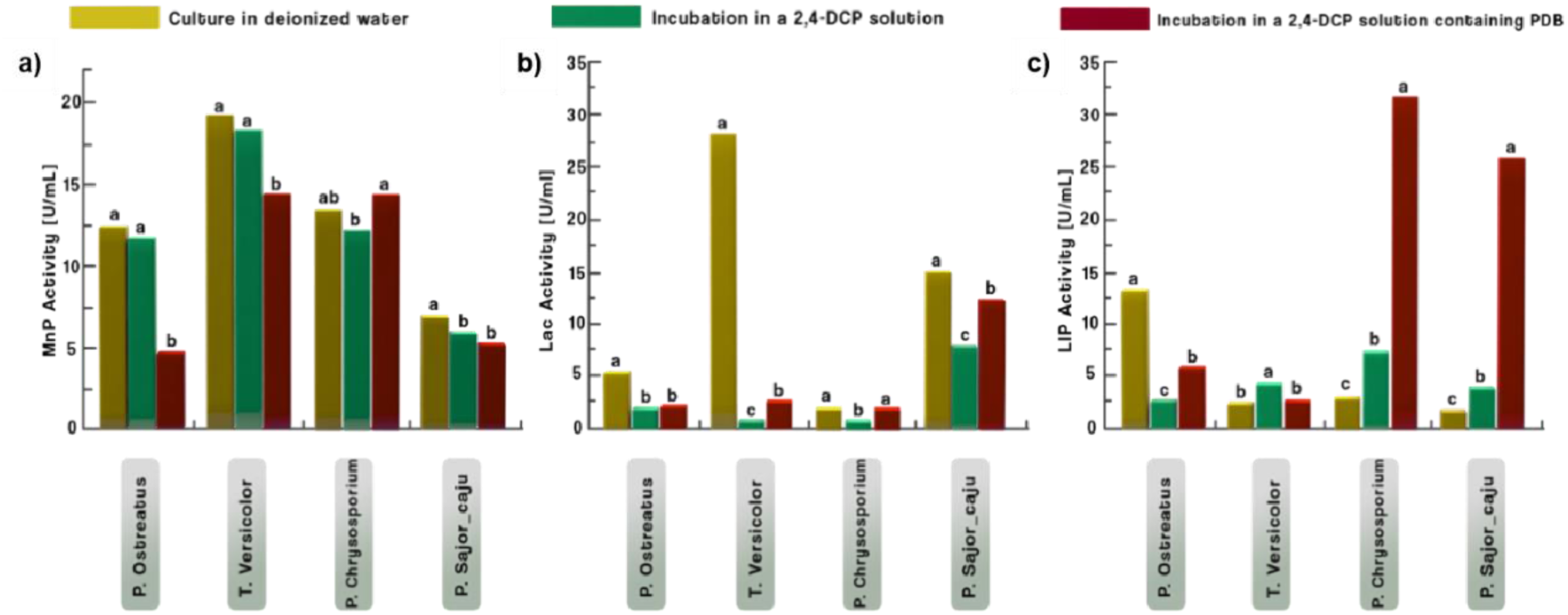
Viability of *MnP*, *Lac* and *LiP* enzymes in each bacterium (a, b, c). Different environments can affect the secretion of extracellular enzymes. Although the enzyme activity of WRF at the beginning of the degradation reaction is inhibited in environments containing only 2,4-DCP, high activity can be maintained in other environments. However, after the "toxic adaptation period", the enzyme activity of white rot fungi increases significantly during the subsequent degradation process because of 2,4-DCP stimulation "toxic excitation" (which can be reacted from the degradation rate change in Fig. 2)

The survival capacity of *MnP* enzyme in *T. versicolor* was 18.49 U/mL, which was 35.61% and 37.46% higher than that of *P. ostreatus* and *P. chrysosporium*, respectively, and 3 times that of *MnP* enzyme in *P. sajor-caju*. These results indicated that *MnP* enzyme had a high viability in *T. versicolor* (**Fig. 1a**) The *Lac* enzyme activity of the bacterium reached 6.20 U/mL, which was 12 times that of *P. chrysosporium* (**Fig. 1b**). This suggests that it has the strongest ability to secrete *Lac* during the degradation of 2,4-DCP. At the same time, the degradation rate of 2.4-DCP could range companying with the activity of *MnP* and *LiP*, thus, delivering *T. versicolor* a good degradation effect on 2,4-DCP. From **Fig. 1c**, it could be obtained that the *LiP* activity of *P. chrysosporium* on 2,4-DCP was 6.45 U/mL, which was 4 times as much as *P. ostreatus* and 9 times as much as *T. versicolor*. This trend was extremely different compared with other two enzymes. All these results indicate that *LiP* enzyme presents a significant higher activity in *P. chrysosporium*. The further comparison between 2,4-DCP solution group and the deionized water control group are implemented, only the *LiP* enzyme’ activity of the *P. chrysosporium*’s three enzymes increase after adding 2,4-DCP in the deionized water.

Besides, the enzyme activity of *Lac* and *MnP* in *P. sajor-caju* is higher, while the enzyme activity of *Lac* is significantly higher than that of *P. ostreatus* and *P. chrysosporium*. The ability of *P. sajor-caju* to degrade 2.4-DCP mainly comes from their unique *MnP* and *LiP*, among which *MnP* plays a major role in the catalytic reaction of phenolic compounds. *MnP* and *LiP* oxidize phenols to form phenoxy radical intermediates, and at the same time perform oxidative dechlorination on 2.4-DCP. Through the joint action of *MnP* and *LiP*, 2,4-DCP can be degraded and attenuated. We hypothesize that the removal of *P. sajor-caju* is mainly achieved through the secretion of *Lac* enzyme to degrade 2,4-DCP and opposite chlorineatoms.

Furthermore, as illustrated in **Fig. 2**, the degradation rate of the four species was notably higher in the group without the addition of PDB. This suggests that the presence of PDB inhibits the degradation of 2,4-DCP by bacteria. It is postulated that this could be attributed to the dechlorination process of 2,4-DCP following the degradation of extracellular enzymes is impeded by the presence of PDB, resulting in the formation of smaller molecular materials that no longer interact with the extracellular enzymes. During this process, *T. versicolor* achieved equilibrium first at 48 hours with a degradation rate of 81.78%, while *P. sajor-caju* exhibited a degradation rate of 78.20% at 120 hours. Combining the experimental findings, it can be concluded that *T. versicolor* and *P. sajor-caju* possess comparative advantages in their ability to degrade 2,4-DCP.

**Fig. 2.**
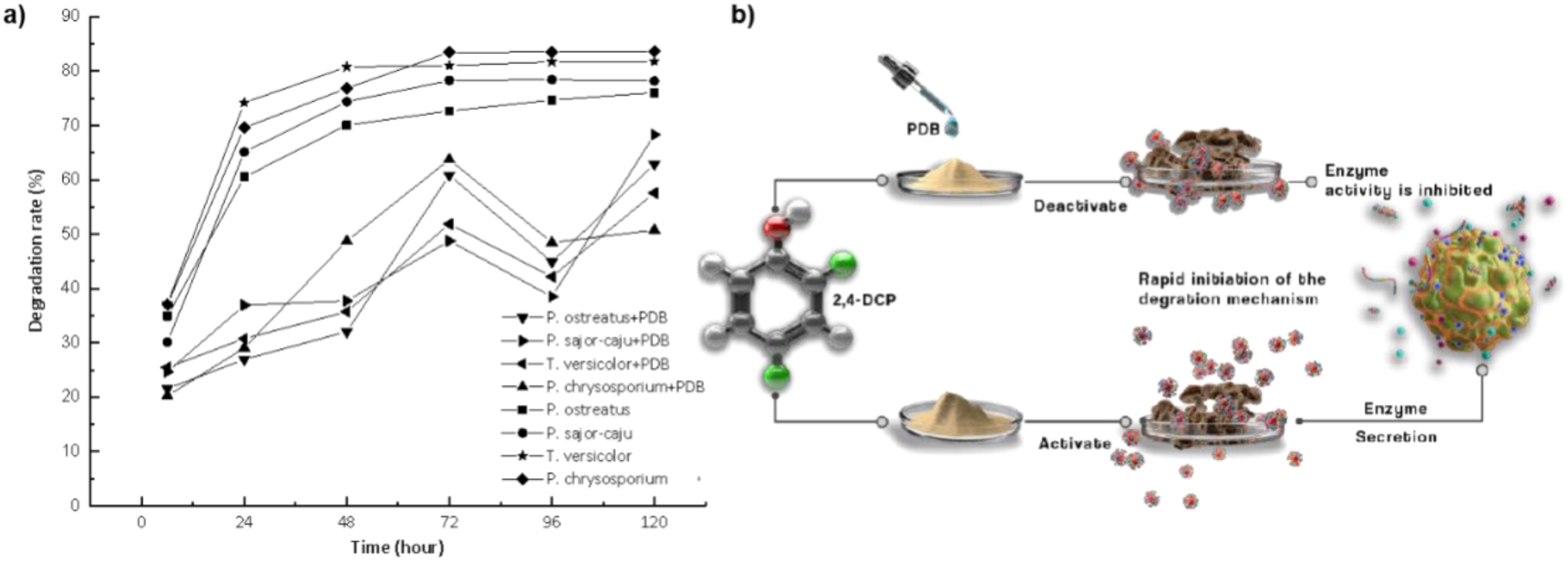
Situation of four WRF under different conditions with decreased solution 2,4-DCP. Degradation of 2,4-DCP by each bacterium. PDB addition experiments demonstrated the extreme ability of WRF to degrade 2,4-DCP in the absence of nutrients. The results of the control test were obtained by the addition of PDB with the no PDB group. According to the experimental results of (a), (b) shows the inhibitory effect of PDB on the degradation function of white rot fungi and the possible activation effect of extracellular enzyme activity when PDB is not added.

### 3.2 Degradation effect of 2,4-DCP under different mixed bacterial groups

The efficiency and ability to degrade pollutants vary depending on the selection of white rot fungi. It is speculated that there may be interactions between the fungi, leading to complementary advantages. To verify this hypothesis, the degradation rates of mixed strains with different groups is compared and the optimal combination is going to be determined. Significant differences in the degradation rate of 2,4-DCP can be observed with different mixed types (See **Table S1**). Some groups achieved final degradation rates exceeding 80%, with the highest degradation rate observed in the group consisting of 1/3 *T. versicolor* and 2/3 *P. sajor-caju* at 84.63±0.28%. When compared to the results of degradation by a single strain, it was evident that the degradation rates of most mixed fungi types were higher than those of single strains due to the synergistic effect between mixed fungi, resulting in an enhanced degradation ability of WRF through interactions.

Furthermore, *P. ostreatus* was found to produce fewer enzymes, leading to its lower contribution to mixed degradation when combined with *T. versicolor* compared to *P. sajor-caju*. However, the results of mixing two and three species of WRF and their symbiotic effects did not exhibit a clear regular pattern. As a result, compatibility tests were conducted on the four fungi shown in **Fig. 3**, no obvious inhibition of the four fungi or transparent bacteriostatic rings were observed. Subsequently, after the two bacteria points were connected to PDA solid medium for four days, both bacteria were found to survive. From the perspective of growth advantages, the growth trend of *T. versicolor*/*P. sajor-caju*, *T. versicolor*/*P. ostreatus*, and *P. ostreatus*/*P. sajor-caju* were similar. When the three strains were mixed, they exhibited normal growth and grew faster. However, when the four strains were mixed, the growth rate of *P. chrysosporium* was notably faster than that of the other strains, potentially leading to nutrient competition in the later stage and inhibiting the growth of the other three strains. Consequently, it was determined that *P. chrysosporium* should be inoculated three days later than the other three fungi in the subsequent experiment due to its poor compatibility with the other three fungi, resulting in generally lower degradation rates in mixed groups containing *P. chrysosporium*. It was preliminarily observed that *T. versicolor* and *P. sajor-caju* exhibit relative advantages over the other two bacteria in degrading chlorophenolic wastewater, although their optimal mixing ratio remains to be explored.

**Fig. 3.**
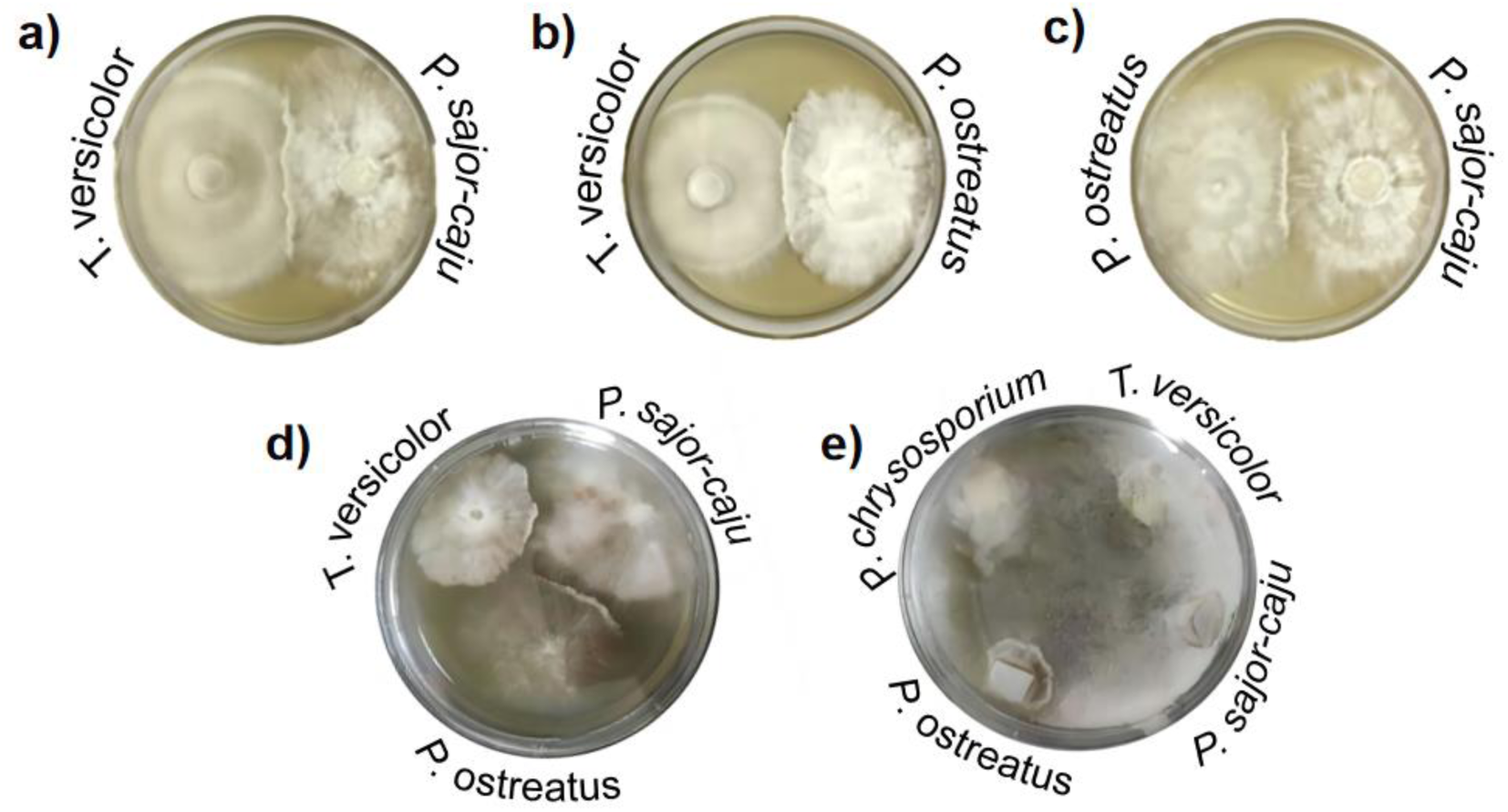
Mixed WRF compatibility test results in PDA. The mixed types in the figure were *T. versicolor*/*P. sajor-caju*, *T. versicolor*/*P. ostreatus*, *P. ostreatus*/*P. sajor-caju*, *T. versicolor*/*P. sajor-caju*/*P. ostreatus*, and four WRF.

The above mentioned results could confirm the effect of different mixed strains proportion to the 2,4-DCP degradation, while detailed composition of the mixed species remains obscure. Hence, JMP software is applied to predict a reasonable model with the initial data. As depicted in **Fig. S1**, the analysis revealed a correlation between the degradation rate and the concentrations of *P. ostreatus* and *P. chrysosporium*, indicating a decrease in degradation rate with increasing content of these strains. Moreover, when the ratio of T. versicolor to P. sajor-caju approached 1:1, the degradation efficiency reached 85.81%. To validate this finding, a parallel experiment was conducted at this ratio as presented in **Table 1**, confirming a final degradation rate of 86%, consistent with the predicted outcome. Hence, this specific ratio is deemed optimal for degrading 2,4-DCP.

**Table 1.** The prediction of the mixing ratio and parallel test of degradation of *T. versicolor* and *P. sajor-caju* with ratio of 1 to 1

| Group | 6 h | 24 h | 48 h | 72 h | 96 h |
| --- | --- | --- | --- | --- | --- |
| 1 | 48.83%±0.71 | 79.74%±0.76 | 82.82%±0.34 | 85.93%±0.23 | 86.09%±0.65 |
| 2 | 47.52%±0.67 | 76.7%±0.26 | 83.77%±0.59 | 84.89%±0.89 | 86.97%±0.56 |
| 3 | 45.31%±0.35 | 80.65%±0.64 | 81.46%±0.45 | 86.67%±0.48 | 86.93%±0.49 |
| 4 | 43.47%±0.56 | 72.83%±0.35 | 82.80%±0.68 | 83.56%±0.61 | 88.33%±0.87 |
| 5 | 47.49%±0.48 | 73.03%±0.46 | 83.87%±0.67 | 85.95%±0.34 | 85.93%±0.45 |

Observing the degradation profile, it is evident that the degradation rate exhibited a notable surge within the initial 24 hours, followed by a gradual tapering off until reaching a state of equilibrium. This phenomenon can be attributed to the accumulation of metabolites, which impeded the physiological and enzymatic activities of the fungi, consequently slowing down the reaction kinetics. Additionally, the decrease in 2,4-DCP concentration within the reaction system may have led to a depletion of carbon sources, thereby diminishing fungal activity. Consequently, addressing issues related to fungal growth conditions and nutrient availability is imperative to achieve optimal degradation of 2,4-DCP.

### 3.3 Preparation and performance determination of immobilized microspheres

Immobilization is a commonly employed technique in microbial experiments, and the immobilized co-culture method plays a crucial role in establishing a stable symbiotic system (Tang et al., 2022). In our endeavor to enhance the degradation efficiency of mixed strains, we adopted an integrated approach involving adsorption, encapsulation, and cross-linking techniques for compound fixation. This method involved the addition of PVA, SA, biochar, SiO_2_, and zeolite powder to a solution of sodium dihydrogen phosphate to create a gel matrix. Within this matrix, the spores of *T. versicolor* and *P. sajor-caju* were immobilized in a 1:1 ratio to form fixed pellets. Notably, the mass concentration ratio of SA to biochar was 1:2. Specific details of the experimental groups are provided in **Table S2**. The 96-hour degradation rate of the immobilized pellets reached an impressive 90.54%, indicating a 4.54% increase compared to the free mixed strains.

The appearance of the immobilized pellets was characterized by a relatively uniform spherical shape, indicating the establishment of a stable and compact symbiotic structure among the encapsulated bacteria (**Fig. 4**). This structure is conducive to bacterial survival and the maintenance of metabolite activity. Further details regarding the properties of the immobilized microspheres can be found in **Table S3**.

**Fig. 4.**
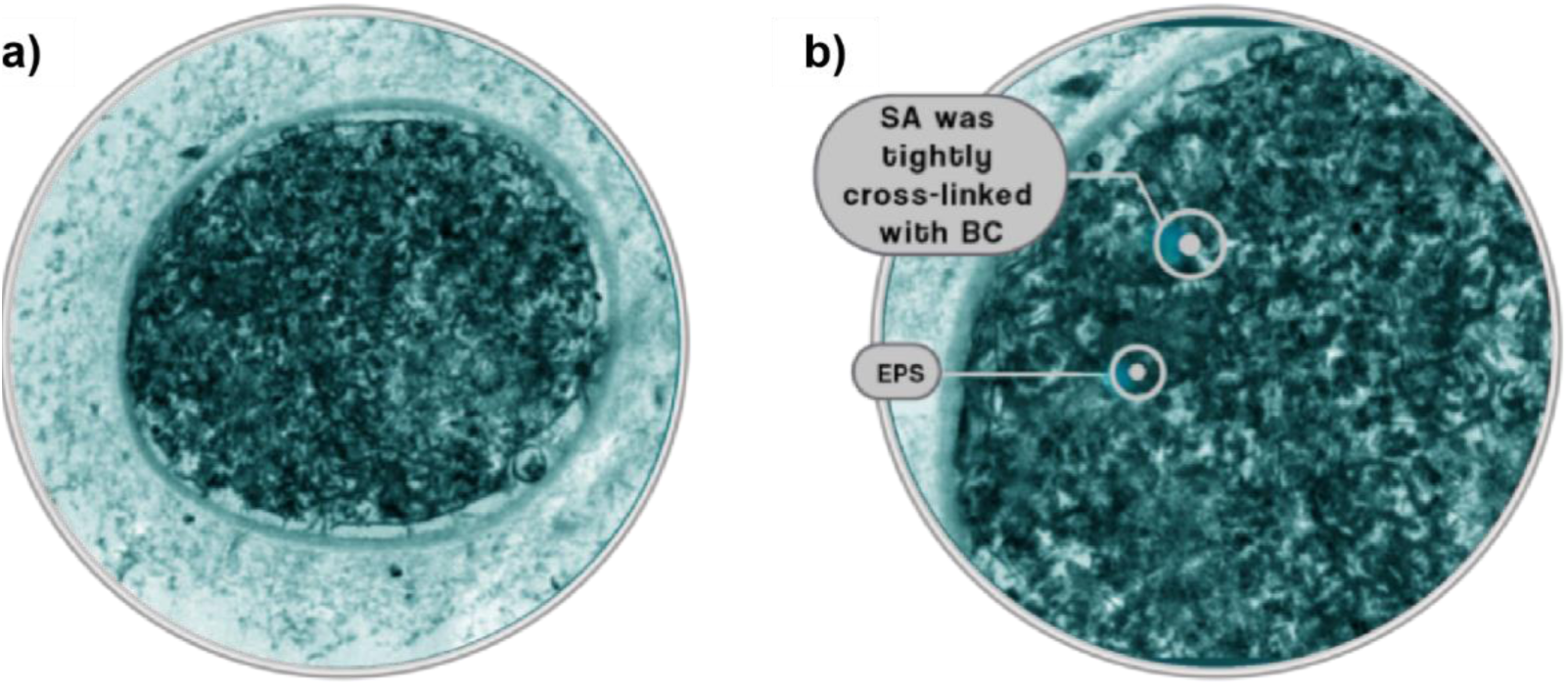
*T. versicolor* and *P. sajor-caju* co-fixed electron microphotographs. After immobilization, the hyphae are tightly wound with each other to form a dense structure. It can be seen in **Fig. 4a** that the shape of the pellet is relatively regular, the black part is the cross-linked hyphae, and the gray outer ring of the pellet is the immobilized material. **Fig. 4b** shows the local EM amplification diagram, which can clearly observe that SA is closely connected to BC, and the EPS secreted by white rot fungi to resist the bad environment.

Regarding the reusability of the globules, analysis presented in **Table S4** demonstrates that upon initial utilization, the PVA pellets exhibited a high degradation rate of 2,4-DCP, achieving approximately 95% after 96 hours with exceptional treatment efficacy. However, with repeated application, the available substances supporting mixed bacterial growth within the PVA pellets were gradually depleted, leading to a decline in bacterial activity and a subsequent weakening of 2,4-DCP degradation capability. Besides, as indicated in **Table S5**, the PVA pellets as prepared remained structurally intact in solution for at least 120 hours, retaining their spherical form for approximately 192 hours thereafter without damage, indicating their robust stability and reusability, rendering them the highly suitability for the treatment of chlorophenol-containing organic wastewater.

### 3.4 Optimization of response surface conditions for degradation of immobilized microspheres

To achieve a higher degradation rate, the influence of environmental factors on the degradation effects are investigated by response surface analysis of single factor combination experiment. Herein, design expert was used to take the response surface analysis and the degradation rates (6 h; 96 h) are selected as the main response values. The response surface optimization curve and contour diagram are shown in **Fig. 5**.

**Fig. 5.**
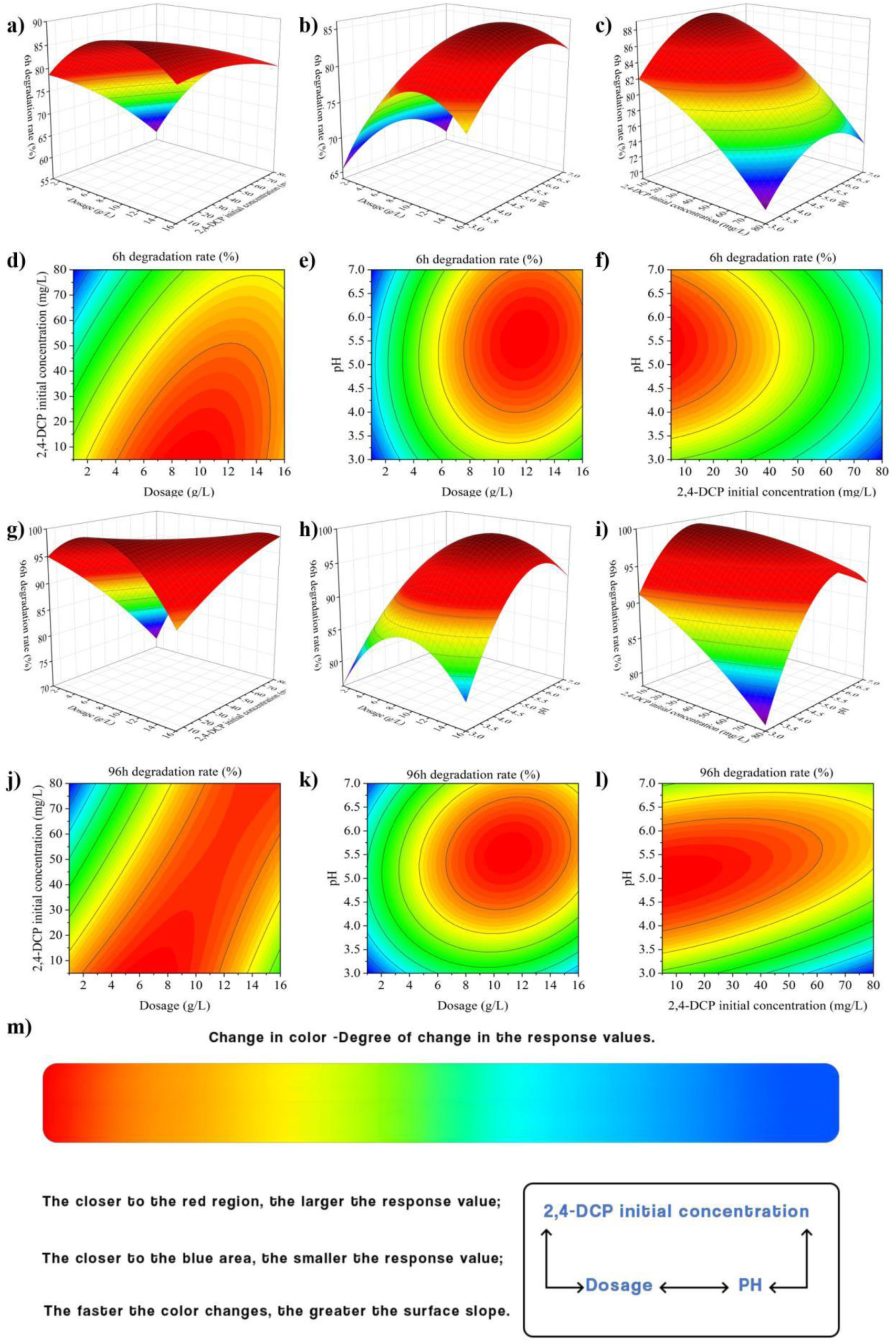
Response surface optimization and contour diagram of immobilized microsphere degradation condition (6 h, 96 h). (a-f) are the results of the experiments with immobilized sphere at 6 h, and (g–l) are the results at 96 h. (m) is the interpretation of the response surface correlation principle.

After comprehensively comparing the effects of the three factors on the degradation rate, the interaction between the environmental pH value and the other two factors was found to be quite significant. In addition, it can be seen from the change of surface inclination and the color change of the corresponding contour map that the main factors affecting the degradation rate of 2,4-DCP are in the following order: concentration of bacteria (g/L) > initial concentration of 2,4-DCP (mg/L) > pH.

**Fig. 6a** is the results of the nine sets of experiments presented in **Table S5**. It is found that the degradation rate increases with the rise of bacteria concentration in this range. However, regardless of different changes in the initial concentration of bacterial dose or 2,4-DCP, the degradation effect always remained optimal at pH 5.5. Besides, as shown in **Fig. 6b**, the degradation effect of 2,4-DCP was the most significant at 96 h under the conditions of bacteria dosage of 11.5 g/L, initial 2,4-DCP concentration of 40 mg/L and pH of 5.5. For this optimal degradation condition, many confirmatory experiments are implemented, and the experimental results showed that the degradation rate of 2,4-DCP could reach (99.07±0.26) % under this condition, which is excellent in practical feasibility.

**Fig. 6.**
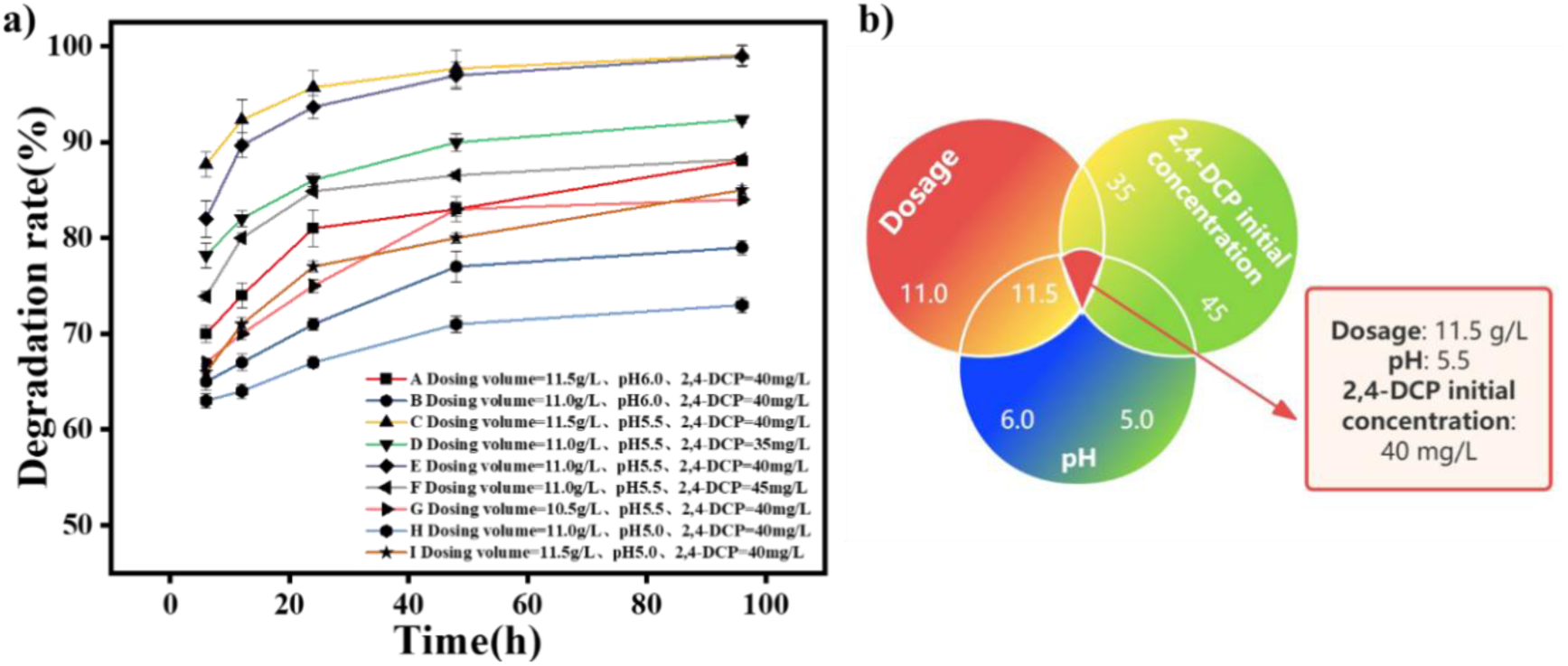
Degradation rate of 2,4-DCP under different conditions of immobilized microspheres. (**a)** is selected by the response surface optimization experiment of 9 groups of degradation rate is relatively high experiment, (**b)** is based on the summary of **(a)** results, the highest degradation rate of the three factors, each area color change of numerical value, different colors in different regions represent different influence, the strongest red, blue influence, the weakest influence size is obtained.

### 3.5 Kinetic curve of 2,4-DCP dechlorination reaction by mixed strains

Since the final degradation products of 2,4-DCP are CO_2_ and H_2_O, this is mainly completed by the dechlorination step, and there is a sequential dichlorination process. Whether this sequence is influenced by the strain species remains obscure. **Fig. 7a** shows the changes of phenolic substances in the system after mixing 2, 4-DCP with a mass concentration of 20 mg/L at a ratio of 1:1. The content of 2, 4-DCP decreased to 0 at 30 h, while the content of 4-CP and 2-CP increased first and followed by decreasing to 0 gradually. In particular, the content of 4-CP reached the highest at 12 h, which was much more than 2-CP. The degradation rate of 2, 4-DCP was the highest at 6-12 h, and the content of 2-CP and phenol increased. Indicating the mixed strain began to secrete *LiP*, *MnP*, and *Lac* to degrade 2, 4-DCP 6 h after adaptation to the system environment and entered the stable stage 12 h later. In addition, the mixed strain achieved complete degradation of phenols within 48 h, and finally achieved complete removal of 2, 4-DCP.

**Fig. 7.**
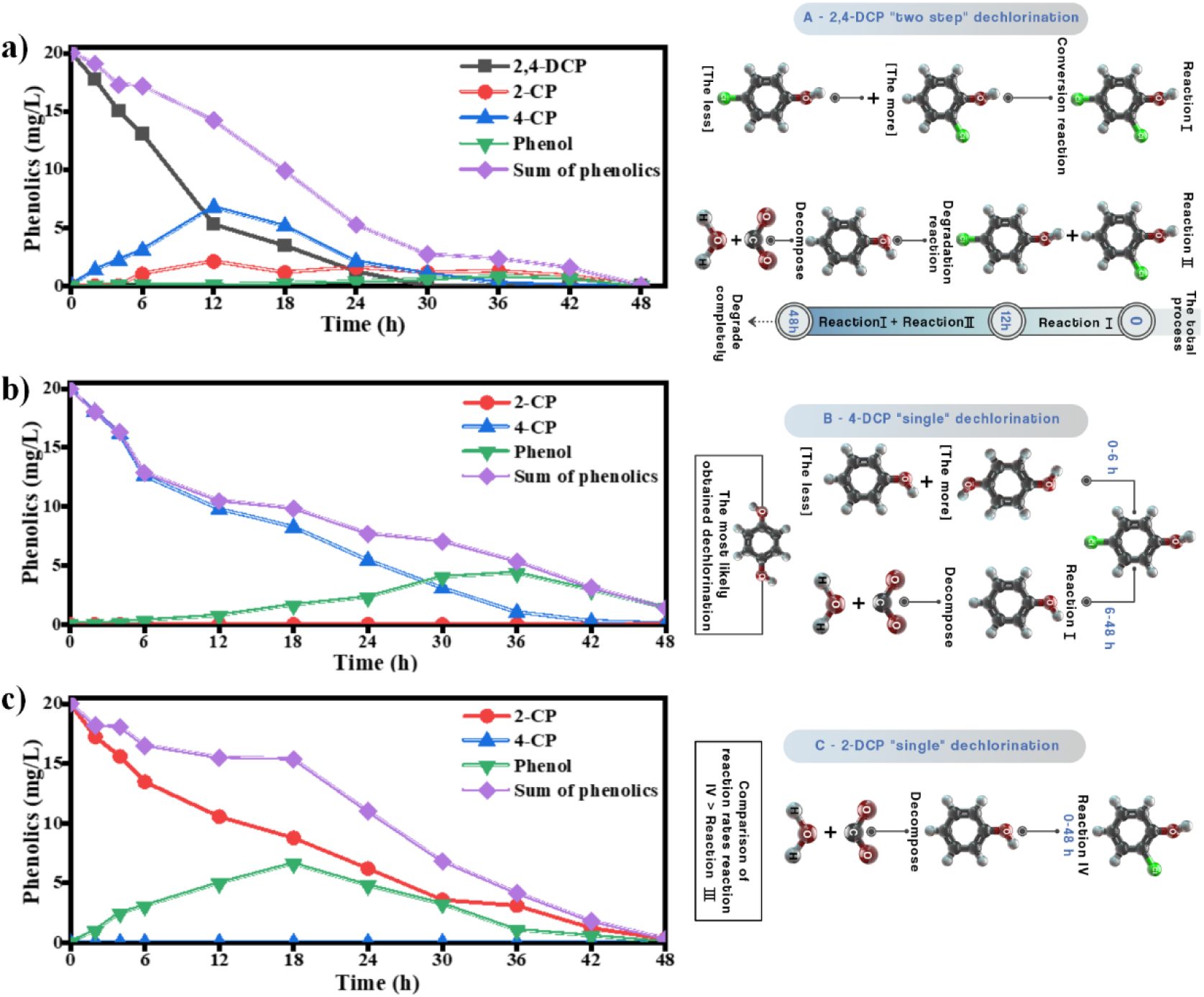
the change of phenolic substances in the system after the degradation of 2,4-DCP with mixed bacteria. (a) describes the process of the gradual dechlorination of 2,4-DCP to produce small-molecule harmless compounds such as carbon dioxide and water. (b) which is the subsequent dechlorination process of 4-CP which is the 2-site dechlorination product of 4-DCP. (c) describes the subsequent dechlorination process of 2-CP which is the 4-site dechlorination product of 2,4-DCP. Mechanism diagram on the right compares the changes in the types and numbers of phenolic compounds in different time systems, describing the reaction process of (a), (b) and (c) in the form of reaction, intuitively obtaining the speed of adjacent and relative dechlorination.

**Fig. 7b** shows the single dehlorination process of 4-CP, it obviously dropped between 0-6 h, but the theoretical degradation product phenol did not rise significantly. It may be resulted from the formation of other phenolic substances, such as hydroquinone (Zhao et al., 2022). The change of 4-CP and phenol was similar at 6-48 h, indicating the dechlorination reaction at position 4 mainly occurred during this period. **Fig. 7c** presents the single dechlorination process of 2-CP, mainly occurred at position 2. In summary, Mechanism diagram on the right of **Fig. 7** indicated the two white rot fungal mixtures would preferentially remove adjacent chlorine atoms on 2, 4-DCP to produce 4-CP. At the same time, they would also choose to remove the chlorine atoms to produce a small amount of 2-CP, and then further dechlorinate to produce phenol. Further, for the degradation of 4-CP and 2-CP, 4-CP was more easily converted into phenol than 2-CP within 0-6 h.

## 4. Discussion

### 4.1 Degradation of 2,4-DCP efficiency decided by co-cultivation technology of WRF

Chlorophenols (CPs) represent a class of hazardous waste predominantly produced during chemical processes. Owing to its extensive application in both manufacturing and agricultural fields, 2,4-DCP has emerged as one of the most common chlorophenols found in aquatic and terrestrial environments globally (Al-Abri et al., 2019). Residues of 2,4-DCP have been consistently detected in soil and water sources across the globe, with residual concentrations varying between 0.02 and 2.2 mg/L. The presence of 2,4-DCP residues poses a significant ecological threat and public health risk due to its high toxicity and persistence in the environments (Kocaoba & Arısoy, 2011). Consequently, the remediation of 2,4-DCP from polluted environment is of paramount importance. The ligninolytic enzyme systems of white rot fungi, including *LiP*, *MnP*, and *Lac*, have been identified as crucial in the degradation of organic pollutants. The catalytic mechanisms and degradation capabilities of chlorophenolic compounds vary among these three enzymes. Notably, Laccase facilitates the removal of chlorophenolic substances, particularly by targeting the ortho and para chlorine atoms of 2,4-DCP (Wang et al., 2004), demonstrating a higher efficiency compared to the dechlorination processes involving *LiP* and *MnP* radicals (Wang et al., 2023b).

There were variations in the enzyme production capacity among the four strains of white rot fungi. *P. sajor-caju* exhibited the highest laccase activity, leading to the most rapid degradation of 2,4-DCP within 24 hours. This suggests that *P. sajor-caju* possesses a robust ability to adjust to the initial concentration of the solution, likely attributed to its strong affinity for forming hydrogen bonds with the numerous hydroxyl compounds generated during the degradation process (Choi et al., 2023). The strong binding of the enzyme’s active site to the substrate enhances the degradation rate. Literatures have indicated that *MnP* can enhance the catalytic capability of *Lac* (Sui et al., 2024). Furthermore, *LiP* can catalyze the initial oxidative dechlorination step in the degradation of chlorinated phenols (Kheirkhah et al., 2020). Achieving an appropriate balance of *LiP*, *MnP*, and *Lac* is crucial for enhancing the degradation rate of chlorophenolic compounds, aligning with previous reports by Schmerling et al. (2022). Additionally, nutrient deprivation can prompt the initiation of extracellular enzyme degradation mechanisms in white rot fungi. The outcomes of this study are consistent with the findings of Beltrán-Flores et al. (2023), indicating that the degradation rate of the four strains was significantly higher in the group control compared to the group with PDB. This phenomenon may be attributed to the response of the bacteria to adverse environmental conditions, where the secretion of the three extracellular enzymes of *Lac*, *LiP*, and *MnP* is triggered only under conditions of carbon or nitrogen source limitation. Their secretion is diminished when nutrient availability is sufficient (Chen et al., 2023).

Co-cultivation technology can harness the potential of fungal biosynthesis and the secretion of metabolically active substances through interspecies interactions, thereby enhancing their environmental resilience (García-Martín et al., 2024). Chen demonstrated that co-culturing of *P. chrysosporium* and *Trichoderma viride* requires a specific inoculation interval to achieve optimal coexistence (Chen et al., 2011), suggesting potential incompatibilities between fungi during co-cultivation. Our experiment yielded similar conclusions under identical culture conditions: the presence of *P. chrysosporium* inhibited the growth of other fungi when co-cultivated with three other species. This inhibition could stem from varying growth rates, nutrient compositions, and the production of intermediate metabolites that impede the growth of other fungi (Keller, 2019).

Besides, both *T. versicolor* and *P. sajor-caju* exhibit the ability to secrete extracellular laccase and intracellular enzymes for metabolizing diverse substrates, making them valuable in degrading complex organic matter (Zhang et al., 2017). Crude enzyme extracts from *P. sajor-caju* have demonstrated efficacy in bisphenol A removal (Omoni et al., 2022). Numerous studies have highlighted the compatibility of *T.* versicolor with other fungi, providing it suitable for co-cultivation (Civzele et al., 2023). However, co-cultivation of *P. sajor-caju* with other fungi remains underexplored. In our experiment, we observed a synergistic effect between these two strains during growth, wherein they stimulated each other, resulting in increased yields of *Lac*, *MnP*, and *LiP*, consequently significantly enhancing the degradation rate of 2,4-DCP. This underscores the feasibility and benefits of co-cultivating *T. versicolor* and *P. sajor-caju*.

During co-cultivation, the ratio of fungi influences their metabolic activity and degradation efficiency (Barbelli-Lopez et al., 2024). Utilizing predictive modeling with JMP software, we observed that when *T. versicolor* and *P. sajor-caju* were mixed at a ratio of 1:1, the degradation rate of 2,4-DCP was maximized. This predicted ratio aligns with the initial inoculation ratio reported by Sun et al. (2023). We conducted experimental validation of the 1:1 ratio of *T. versicolor* and *P. sajor-caju*, confirming that at this ratio, the two fungi effectively distributed nutrients, established a symbiotic relationship, and achieved the highest degradation rate of 2,4-DCP at 86%, surpassing other mixed ratios. In experiments with alternative proportions, we observed that when the proportion of one fungus inoculated exceeded 0.5, the fungus at the higher proportion gained a growth advantage, rapidly depleting carbon sources and inhibiting the growth of the other fungus, consequently reducing the degradation rate significantly (Gao et al., 2024).

### 4.2 Immobilized WRF microbial remediation with enhanced stability

Natural cells, especially prokaryotic cells, tend to move and absorb nutrients in their microenvironment (Zhang et al., 2019). Compared to free microorganisms, immobilized microorganisms exhibit greater adaptability and tolerance to toxic substances, making them suitable for water and soil environment treatment (Rong et al., 2024). It was observed that the enzyme activity of immobilized white rot fungi began to surpass that of free white rot fungi after the fifth day of culture, with significantly higher activity observed at later stages. Similar conclusions were drawn from this experiment, demonstrating the enhanced adaptability of immobilized white rot fungi to oscillatory culture. This adaptability stems from their ability to minimize mycelium damage caused by shear force, effectively enhancing the stability and enzyme activity of white rot fungi, and facilitating the continuous degradation of 2,4-DCP pollutants in wastewater. Moreover, the PVA, SA, and BC utilized in this study are environmentally benign, serving solely to embed microorganisms and absorb harmful substances without desorbing over time. Therefore, the utilization of biochar-sodium alginate fixation technology is crucial for improving the tolerance and degradation efficiency of degrading bacteria.

Second, the quantity of immobilized white rot fungi is pivotal in the actual degradation process of 2,4-DCP. In this study, the optimal concentration was determined to be 11.5 g/L. Beyond this threshold, enzyme activity diminished and the degradation efficiency declined. This reduction is likely due to the aerobic nature of white rot fungi, coupled with a possible scarcity of oxygen in the solution. Moreover, an excess of PVA microspheres can lead to aggregation, diminishing the effective contact area with the wastewater. A rapid consumption of nutrients, leading to competition, is another significant factor. These findings suggest that an optimal bacterial count is necessary to trigger secondary metabolism and progress to the biodegradation phase. The initial concentration of the 2,4-DCP solution also influences the degradation rate. Shi et al. (2024) observed that the initial concentration of wastewater facilitated the treatment by white rot fungi at low levels but was inhibitory at higher concentrations. This research revealed that immobilized white rot fungi exhibit hormesis in treating 2,4-DCP. At low concentrations, the fungi’s high-affinity receptor subtypes are activated, enabling the organism to stabilize and develop tolerance. Conversely, in environments with high 2,4-DCP concentrations, the activity of the lignin oxidase system is suppressed, adversely impacting the degradation efficiency.

Additionally, alongside concentration and dosage, pH stands out as a critical external factor impacting the degradation efficacy of fungal strains. This study reveals that an acidic milieu fosters white rot fungi growth, with the optimal pH identified as 5.5. Conversely, relatively high pH levels exhibit a certain inhibitory effect on white rot fungi growth. Within the pH range of 5.0-5.5, immobilized microspheres maintain a commendable degradation rate, indicating the robust tolerance of white rot fungi under such conditions, aligning closely with the findings of Jeganathan et al. (2024).

Moreover, Velásquez-Quintero et al. (2022) highlight the interactive role of dosage and initial organic concentration in both biological and chemical degradation processes. In this experiment, the response surface optimization method was employed to conduct intuitive and range analyses among influencing factors, yielding similar results. Notably, dosing amount and pH exhibit significant interaction. When they operate together, their impact on degradation rates under individual factors can be notably attenuated. Through an appropriate balance, the inhibitory effect of specific surface area of immobilized pellets and the toxicity of chlorophenols on mixed bacteria can also be effectively mitigated.

### 4.3 Detailed mechanisms of the 2,4-DCP degradation with the solid mixed strains microsphere agent

Generally, CPs exhibit significantly lower biochemical viability compared to typical aromatic compounds due to molecular structural alterations induced by chlorine atoms. Dechlorination represents a pivotal stage in CPs’ biodegradation (Li et al., 2023), with dechlorination reactions predominantly governing the biodegradation of chlorinated organic pollutants. Elaborate mechanisms are elucidated in **Scheme 1a** (Quan et al., 2003). Upon determining the appropriate dosing quantity, initial 2,4-DCP concentration, and pH for a 1:1 mixture encapsulation of *T. versicolor* and *P. sajor-caju*, we established a kinetic model for 2,4-DCP dichlorination. Our findings indicate that: at elevated CPs concentrations, the mixed bacteria predominantly catalyze dechlorination at position 2, yielding a substantial quantity of phenol; conversely, at lower CPs concentrations, dechlorination primarily occurs at position 4, producing minimal 2-CP dechlorination and subsequent phenol formation. Consequently, we infer the subsequent degradation process of 2,4-DCP by mixed strains:

**Scheme. 1.**
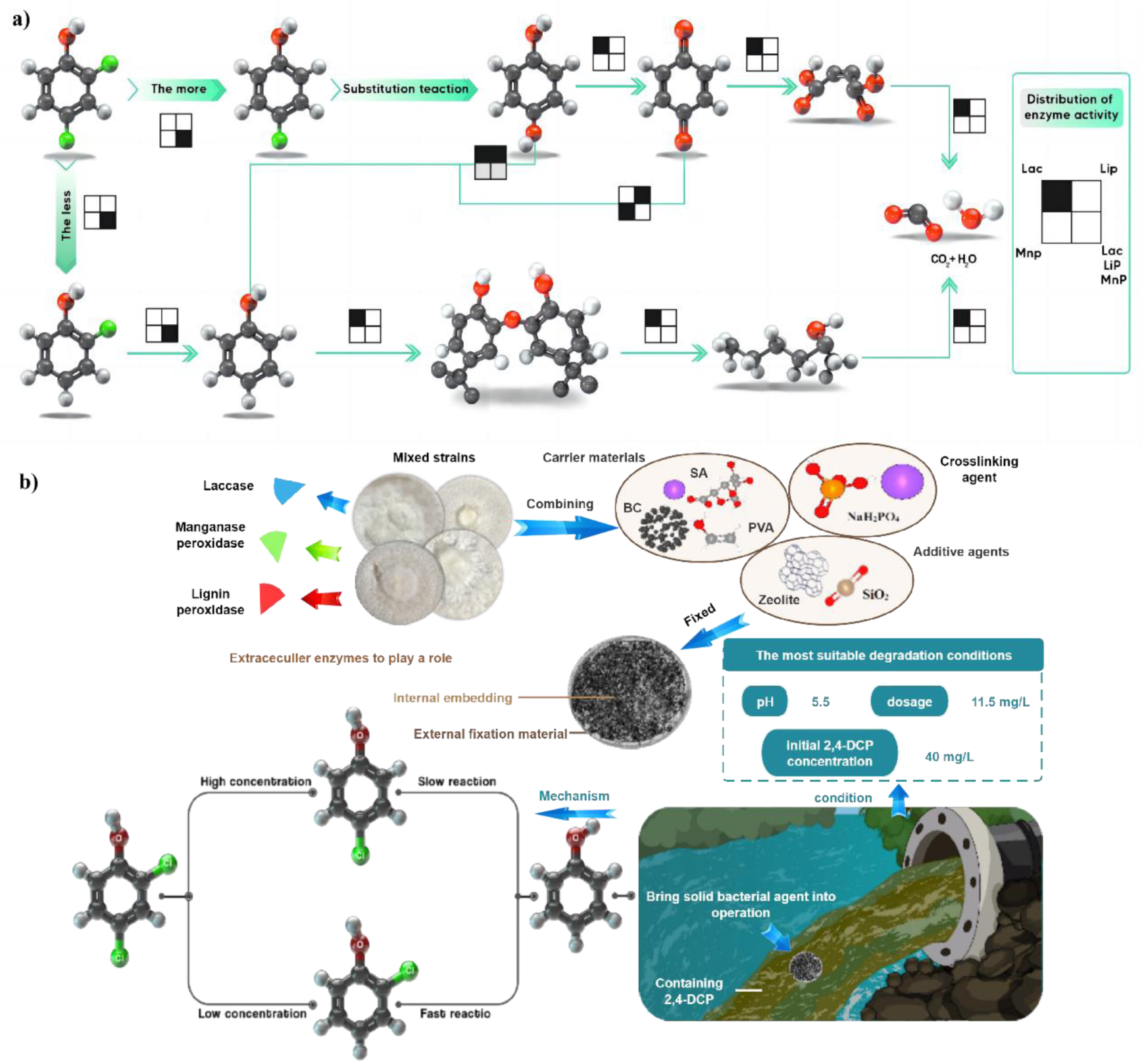
Degradation mechanism of 2,4-DCP. (a) is a summary of the total degradation process of 2,4-DCP, including the extracellular enzymes involved in each reaction stage, based on the research results of other researchers. (b) is a summary of the process of degradation of 2,4-DCP by mixed strains, presenting to the synergistic mechanism of co-cultivation of WRFs.

Initially, o-chlorophenol undergoes faster degradation than p-chlorophenol under laccase catalysis, likely attributable to the activation of the benzene ring by reactive groups such as -OH, facilitating rapid reaction with electrophilic substituents (Eastwood et al., 2011). Given that 2,4-DCP presents two chlorine atom substitution positions, namely position 2 and position 4, *Lac* catalyzes dechlorination at these sites initially, with a propensity for dechlorination at position 2. At heightened concentrations, a greater proportion of 2,4-DCP undergoes dechlorination to form 4-CP, with a minor fraction dechlorinating to form 2-CP. Subsequently, in the presence of *Lac*, both 4-CP and 2-CP proceed to produce phenol. Notably, the subsequent phenol production from 4-CP and 2-CP varies; 2-CP undergoes rapid conversion to phenol for degradation, whereas 4-CP exhibits slower dechlorination, leading to a diminished degradation rate. Furthermore, phenol levels decrease in later reaction stages, possibly due to continued catalytic oxidation by *Lac*, yielding active quinones that polymerize and mitigate toxicity (Yoon et al., 2022). With the synergistic cooperation of the mixed cultivation of different WRF, the chlorophenol-contaminated wastewater could be degraded efficiently and intelligently (**Scheme 1b**).

## 5. Conclusions

In response to chlorophenol-contaminated wastewater, a pioneering approach involves the strategic selection of diverse strains, optimizing the preparation of immobilized pellets, and fine-tuning environmental conditions is investigated. The synergy among mixed strains facilitates the establishment of a more efficient enzyme cascade reaction, enhancing mass transfer while ensuring microsphere stability through the reasonable immobilization strategy. The resulting PVA-immobilized mixed bacteria exhibit substantial 2,4-DCP degradation, laying a solid foundation for further research into fermentation, immobilization formulations, and the utilization of WRF, thereby promoting both economic and ecological benefits. Simultaneously, exploration of the dechlorination pathway for chlorophenol pollutants typified by 2,4-DCP offers insights to enhance treatment methodologies for chlorophenol-contaminated environments.

## List of abbreviations

2,4-DCP: 2,4-dichlorophenol
CPs: Chlorophenols
DAD: diode array detector
HPLC: high-performance liquid chromatograph
*Lac*: *laccase*
*Lip*: *lignin peroxidase*
*Mnp*: *manganese peroxidase*
PAHs: polycyclic aromatic hydrocarbons
PCBs: polychlorinated biphenyls
PCDDs: polychlorinated dibenzo-p-dioxins
PVA: polyvinyl alcohol
ROS: reactive oxygen species
SA: sodium alginate
WRF: White Rot Fungi

## Acknowledgements

This study was supported by the National Natural Science Foundation of China (32001224 and 41702181); and the University Students Innovation and Entrepreneurship Training Program of Zhejiang Province of China (202310345048). Y. Orooji acknowledges the financial supports rendered by Natural Science Foundation of Zhejiang Province (Nos. LD21E080001), and Zhejiang Provincial Ten Thousand Talent Program (ZJWR0302055).

## Author Contributions

P.L. and Y.O. conceived the work. H.L., X.L., Y.S designed the experiments. H.L. and W.X. wrote the manuscript and finalized the work. Z.W., J.X., K.C., D.C., and Q.X. participate this work. F.F. provided helpful discussions and all authors discussed the experimental results.

## Appendix A. Supplementary data

Supplementary information is available for this paper. Including the **Fig. S1** and Table S1∼S6.

